# In vivo mapping of protein-protein interactions of schizophrenia risk factors generates an interconnected disease network

**DOI:** 10.1101/2023.12.12.571320

**Authors:** Daniel B. McClatchy, Susan B. Powell, John R. Yates

## Abstract

Genetic analyses of Schizophrenia (SCZ) patients have identified thousands of risk factors. In silico protein-protein interaction (PPI) network analysis has provided strong evidence that disrupted PPI networks underlie SCZ pathogenesis. In this study, we performed *in vivo* PPI analysis of several SCZ risk factors in the rodent brain. Using endogenous antibody immunoprecipitations coupled to mass spectrometry (MS) analysis, we constructed a SCZ network comprising 1612 unique PPI with a 5% FDR. Over 90% of the PPI were novel, reflecting the lack of previous PPI MS studies in brain tissue. Our SCZ PPI network was enriched with known SCZ risk factors, which supports the hypothesis that an accumulation of disturbances in selected PPI networks underlies SCZ. We used Stable Isotope Labeling in Mammals (SILAM) to quantitate phencyclidine (PCP) perturbations in the SCZ network and found that PCP weakened most PPI but also led to some enhanced or new PPI. These findings demonstrate that quantitating PPI in perturbed biological states can reveal alterations to network biology.

## Introduction

Understanding how proteins interact to form networks is essential to understanding human disease pathogenesis[1]. A single genetic coding mutation can disrupt an entire protein-protein interaction (PPI) network or create a disease specific network[2]. Since proteins form associations with multiple proteins, a single gene mutation could perturb numerous PPI networks, which may be why one mutation can underlie multiple disease phenotypes. In diseases with multiple genetic causes or risk factors, gene products associated with the same or similar diseases tend to interact with each other[3–5]. It is posited that multiple genes conferring low to moderate vulnerability to a disease individually can produce more significant disease potential when they are localized within the same PPI network[6]. Understanding what networks are disrupted in a disease state could result in the identification of mechanistic details for the disease.

Analysis of PPI holds great promise for understanding the pathogenesis of schizophrenia (SCZ). SCZ is a chronic debilitating disease which manifests as positive symptoms(i.e. hallucinations and delusions), negative symptoms(i.e. emotional withdrawal and social interaction deficits) and cognitive dysfunction[7]. SCZ patients have difficulty successfully integrating into society, an increased risk of suicide, and a reduction in life expectancy[8–10]. It is estimated that the prevalence of SCZ in most populations[11] is ∼1%. Drug treatment of SCZ has not significantly improved since the development of antipsychotics in the 1950s and antipsychotics only alleviate positive symptoms. The dearth of pharmacological treatments highlights the lack of understanding of the mechanisms behind the pathogenesis of this disease [12]. SCZ is a disease of high heritability, but the complex polygenicity among different populations combined with the involvement of environmental factors has been a challenge to finding drug treatments and cures. About 6000 genes that have been linked to SCZ pathogenesis with only ∼9% assigned a causal relationship[13]. This large number of risk factors may stem from the genetically diverse background of SCZ patients and the heterogeneity of SCZ phenotype. There are a few rare genetic risk factors that individually confer a relatively high risk, whereas more common risk factors (which may require additional environmental risk factors) are postulated to act in an additive manner to trigger disease [14]. PPI disease network analyses have been employed to understand this immensely heterogenous genetic data and have demonstrated that SCZ risk genes interact more significantly with other SCZ risk genes than with genes with no SCZ association[15–17]. These analyses have been able identify previously unknown risk factors that are integral components of enriched SCZ relevant pathways [13, 17]. SCZ risk factors and their PPI networks have been shown to be enriched in synaptic proteins, consistent with the biological evidence that synaptic dysregulation is integral to the pathogenesis[15, 17–20]. There is great optimism for future treatments now that *in silico* PPI analyses have begun to unravel the complex genetic landscape of SCZ.

*In silico* PPI analysis has been hindered by the need for *a priori* knowledge of a gene’s function [21, 22]. Tissue-specific networks have been shown to be more useful for dissecting pathogenesis, since most human diseases are tissue-specific[23]. However, over the last decade, mass spectrometry (MS) has become more widely used for the elucidation of PPI. Using tagged proteins and heterologous expression in cultured cells, thousands of novel PPI have been reported using MS[24–27]. These herculean studies have provided important biological insights, but this data often does not accurately reflect the uniqueness and complexity of brain tissue. Heterologous expression and proteins with ectopic tags also have been reported to alter the native characteristics of endogenous proteins which could affect their PPI[28, 29]. Immunoprecipitation (IP) of endogenous proteins with antibodies (Ab) has also been used to identify PPI. Although an endogenous IP is not as high throughput as a workflow using epitope tagged baits in cell culture, it does provide for PPI in tissue and ultimately animal models of disease [30, 31]. For example, a recent IP-MS study using an Ab to the Akt kinase discovered a novel functional interaction between the Akt and potassium/sodium hyperpolarization-activated cyclic nucleotide-gated channel (HCN1) in the rat hippocampus [32]. This interaction was not identified in two recent large scale PPI datasets in cultured cells using tagged Akt as a bait, suggesting that it is a brain specific interaction [25, 33]. As signaling pathways react to cellular perturbations, PPI also change to support the dynamic milieu. Thus, quantitation of PPI between different physiological states is needed to understand the differences in signal transduction under normal and diseased conditions. PPIs are emerging as attractive targets for drug development[34–36], yet reports of accurate quantitation of IP-MS datasets between disease states in tissue are scarce.

This study sought to construct a brain specific SCZ PPI network by identifying the PPI of multiple proteins implicated in SCZ using the IP-MS strategy. Eight SCZ risk factors were immunoprecipitated from the rat hippocampus, a brain region vulnerable to SCZ pathology[37, 38]. One hypothesis of the pathogenesis of SCZ is the hypofunction of glutamate[39, 40]. This hypothesis is supported by the finding that the synaptic network enriched from SCZ risk genes is intimately connected to N-methyl-D-aspartate glutamate receptors (NMDAR) function[18–20, 41, 42]. Phencyclidine (PCP), a non-competitive inhibitor of the NMDAR, induces SCZ symptoms in humans and exacerbates symptoms in SCZ patients[43]. Animals treated with PCP have been used as models to dissect the neurobiology of SCZ and successfully screen for SCZ drugs [44, 45]. To gain insight into SCZ pathogenesis, we quantitated the effect of PCP on our PPI network.

## Experimental Procedures

### PCP treatment of rats

Sprague–Dawley male rats (Harlan Laboratories, San Diego, CA, USA) weighing 300– 400 g were injected with either saline or PCP (1.25 mg kg^−1^; Sigma Aldrich, St. Louis, MO, USA) at a volume of 1ml kg^−1^. After 26 min, rats were lightly anesthetized with isoflurane and euthanized by decapitation. Brains were removed and hippocampi were dissected, placed in dry ice-cooled isopentane for 5 s, and stored at –80° C. Animals were housed in pairs in clear plastic cages located inside a temperature- and humidity-controlled animal colony and were maintained on a reversed day–night cycle (lights on from 1900 to 0700 hours). Food (Harlan Teklad, Madison, WI, USA) and water were available continuously.

### Metabolic 15N labeling of rat brains

Sprague–Dawley rats were labeled with ^15^N as previously described [46]. At p45 the labeled rats were euthanized using halothane and the brains were quickly removed and then frozen with liquid nitrogen. Eight ^15^N-labeled whole brains were homogenized together in 4mM Hepes pH 7.5, 200mM NaCl with protease and phosphatase inhibitors (Roche, Indianapolis, IN, USA). The ^15^N-labeling efficiency was determined to be 95% using a previously described method [47].

### Immunoprecipitations

Each hippocampus was homogenized on ice in 4mM HEPES, pH7, 200mM NaCl with protease and phosphatase inhibitors with an Eppendorf dounce grinder. Immunoprecipitation (IP) buffer (4mM HEPES, pH7, 200mM NaCl, 0.5% Triton, 0.5% NP-40, and 0.01% Sodium Deoyxcholate) was added to the homogenates and incubated overnight with gentle rotation. The homogenates were centrifuged for 30 min at 17,000 x g. The pellets were homogenized with IP buffer and incubated for 1hour while gently rotating. The homogenates were centrifuged for 30 min at 17,000 x g. The two supernatants were combined, and a BCA protein assay (Fisher Scientific) was performed. Normal sera bound to Protein A (Life Technologies) or Protein G (Abcam) beads were added and incubated overnight to preclear the supernatant. The precleared supernatant was divided into two Eppendorf tubes and beads were either cross-linked to the bait antibody (Grin2b-rabbit #06-600 Millipore,Ppp1ca-Abcam Rabbit #ab16446, Syngap -rabbit #ab3344, Stx1a – Abcam rabbit #ab170890, Map2k1 – Abcam rabbit #ab32091, Syt1 – Abcam mouse #ab13259, Grm5 – Abcam mouse #ab134271, Gsk3b – Abcam mouse #ab2602) or cross-linked to the normal sera of the same species that the bait was added to. Mouse antibodies were crosslinked to Protein G beads and rabbit antibodies were crosslinked to Protein A beads. The beads were incubated with supernatants overnight and then washed three times with 4mM HEPES, pH7, 200mM NaCl. The beads were frozen at -80C until protein digestion. All procedures were performed on ice or at 4°C.

### Protein Digestion

Proteins were eluted from the beads twice with 5% SDS then heated at 90°C for 10 min. The eluate was precipitated with methanol and chloroform. The precipitate was digested with trypsin and ProteaseMAX (Promega) as previously described [48].

### Mass Spectrometry Analysis

Each digestion was analyzed twice by mass spectrometry to create two technical replicates per IP. For each technical replicate, 15ul of the digestion mixture was injected directly onto a 50cm 100um i.d. capillary containing 1.7μm BEH C18 resin (Waters). Peptides were separated at flow rate of 300nl/min on Easy-nLC 1000 (ThermoFisher) on 220 min reversed-phase gradient. Peptides were eluted from the tip of the column and nano electrosprayed directly into an Elite or Velos Pro mass spectrometer (ThermoFisher) by application of 2.5kV voltage at the back of the column. The Grin2b IP was analyzed by MudPIT as previously described[49]. The mass spectrometer was operated in a data dependent mode. Full MS1 scans were collected in the orbitrap at 240K resolution and 20 rapid CID MS/MS in the ion trap [50]. Dynamic exclusion was used with an exclusion duration of 30 sec.

### Interpretation and Quantitation of MS data

Data from technical replicates were combined prior to database searching. MS2 (tandem mass spectra) were extracted from the XCalibur data system format (.RAW) into MS2 format using RawConverter [51]. The MS2 files were interpreted by ProLuCID and results were filtered, sorted, and displayed using the DTASelect 2 program that employed a reversed-sequence decoy database strategy and filtering with a 5 ppm mass accuracy [52, 53]. Searches were performed against UniProt rat database. For each MS dataset (i.e. two technical replicates), the protein false discovery rate was < 1%. The ^14^N/^15^N quantitation and statistical analysis was performed by Census, as previously described [48]. Before statistical analysis, the ^14^N/^15^N protein ratios were normalized by the ^14^N/^15^N protein ratio of the bait in each MS-IP dataset.

### Bioinformatic Analysis

Protein interactors for each bait were identified with SAINTexpress using default values with each SAINT dataset generated from 3 bait IP and 3 control IP[54]. Enrichment of GO terms was performed by WebGestalt [55]. Three separate enrichment analyses were performed for biological function, molecular function, and cellular component (i.e., localization). For all analyses, the FDR was < 0.0001. Some significantly enriched GO terms were combined when shared proteins were greater than 80%. Proteins expressed in the whole brain were used as background to calculate the enrichment. The background brain dataset was obtained from SynGo [56]. To determine if the interactors identified by SAINT have been reported as SCZ susceptibility genes, multiple databases were used that combine datasets from published reports. SCZ GWAS database (ID# MONDO_0005090) was downloaded from the NHGRI-EBI Catalog of human genome-wide association studies. *De novo* mutation datasets and datasets of genes differentially methylated in SCZ were downloaded from the SZDB2.0 [57]. All data was downloaded in March 2022.

## Results

### Selection of SCZ susceptibility genes

Our goal was to identify the PPI networks of SCZ susceptibility genes *in vivo*. The comprehensive SCZ susceptibility gene databases (i.e. SZDB2.0[57] and SZGR2[58]) have integrated published studies analyzing large SCZ cohorts employing a variety of genetic methods, including GWAS, de novo mutation analysis, and differential methylation analysis which have implicated thousands of genes in SCZ. We chose proteins with a synaptic localization for this study since synaptic dysfunction is a central component of SCZ pathophysiology[41]. In addition, we examined phosphorylation changes in SCZ proteins that were induced by PCP [48] since phosphorylation is required for many PPI[59]. However, the major limiting factor of our study was the identification of commercial antibodies capable of IP-MS. In total, eight SCZ susceptibility factors (i.e. Grin2B, Grm5, Gsk3b, Map2k1, Ppp1ca, Stx1a, Syngap1, and Syt1) were employed as baits to identify SCZ PPI networks (**Figure 1A**). In addition to evidence from genetic studies implicating these genes with SCZ, several laboratories have published reports on these baits providing evidence of a causal relationship with SCZ pathogenesis (**Figure 1B**). For example, Grin2b is a subunit of the NMDAR, whose dysfunction is central to the glutamate hypofunction SCZ hypothesis. The baits reflected the functional diversity of SCZ risk factors and included kinases (i.e., Gsk3b and Map2k1), phosphatase (i.e., Ppp1ca), GPCR (i.e., Grm5), ion channel (i.e., Grin2b), GTPase activating protein (i.e., Syngap1), and membrane fusion (i.e., Stx1a and Syt1). These proteins are interconnected in many ways beyond their potential involvement in SCZ pathogenesis. Although most of the baits are localized to multiple subcellular compartments, all proteins have been localized to the synapse [56]. Furthermore, physical and functional interactions have been reported between these baits. For example, Grin2b, Grm5, Map2k1, and Gsk3b are listed as PPI of Syngap1 in BioGrid (The Biological General Repository for Interaction Datasets) database [60]. Functional interactions of the baits include the regulation of Grm5 activation by NMDAR [61].

**Figure 1.**
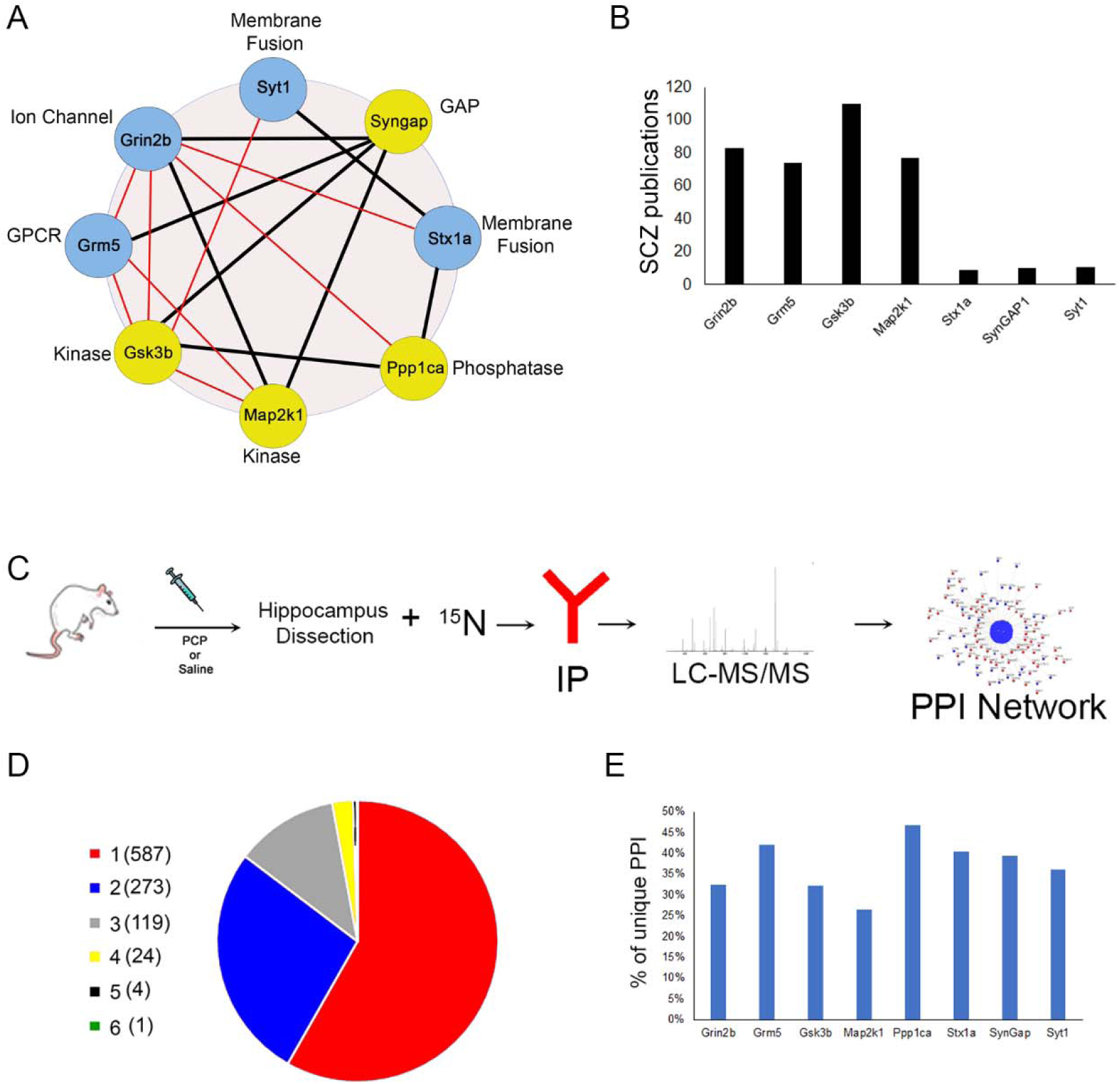
**A**, SCZ susceptibility proteins (i.e., baits) were analyzed. Blue circles represent a transmembrane protein and yellow circles represent a soluble protein. Black lines indicate a physical protein interaction described in the BioGrid database and red lines indicate a functional interaction described in the literature. **B**, The number of publication in Pubmed (6/2023) using gene name and “schizophrenia” as search terms. **C**, Workflow used to generate the SCZ PPI network. **D**, The number of baits assigned to an interactor. **E**, The percentage of unique interactors assigned to each bait network.

### Novelty of the SCZ PPI Network

Figure 1C depicts the experimental workflow. The rats were injected with 1.25mg/kg of PCP or saline (SAL) and were sacrificed after 25min. The hippocampi were dissected. These unlabeled or ^14^N hippocampi were mixed 1:1(wt/wt) with solubilized ^15^N labeled rat brain to serve as an internal standard for the quantitation between PCP and SAL interactomes using MS[46]. The endogenous baits were immunoprecipitated (IP) with antibodies from three PCP treated rats and three SAL injected rats. In parallel, six control IPs (i.e., 3 PCP and 3 SAL) were performed. In total, we performed MS analysis on 96 (48 bait and 48 control) IPs. Protein identifications were similar between the PCP and SAL samples but varied between the different baits (***Supplementary Figure 1A and B***). The baits were enriched at least 9-fold compared to control IPs and there were no significant differences in the baits abundance between SAL and PCP treated brains (***Supplementary Figure 1C***). The proteins identified in the bait IPs were analyzed by the computational tool SAINT (Significance Analysis of Interactome), which determines the probability of a protein interacting with the bait by comparing its abundance distribution to the control IP[62]. This analysis was performed on the ^14^N identifications, and PCP and SAL datasets were analyzed separately. Using a 5% FDR, 1007 unique proteins were deemed to be highly confident interactors. Some proteins were found to interact with multiple baits, but the majority (58%) were specific to one (Figure 1E **& F** and ***Supplementary Table1 & 2***). Overall, 1612 unique PPI were observed. This PPI dataset was compared to PPI deposited in the BioGrid database [60]. All baits had interactors that were verified by BioGrid, but 92% of our PPI were novel (Figure 2A).

**Figure 2.**
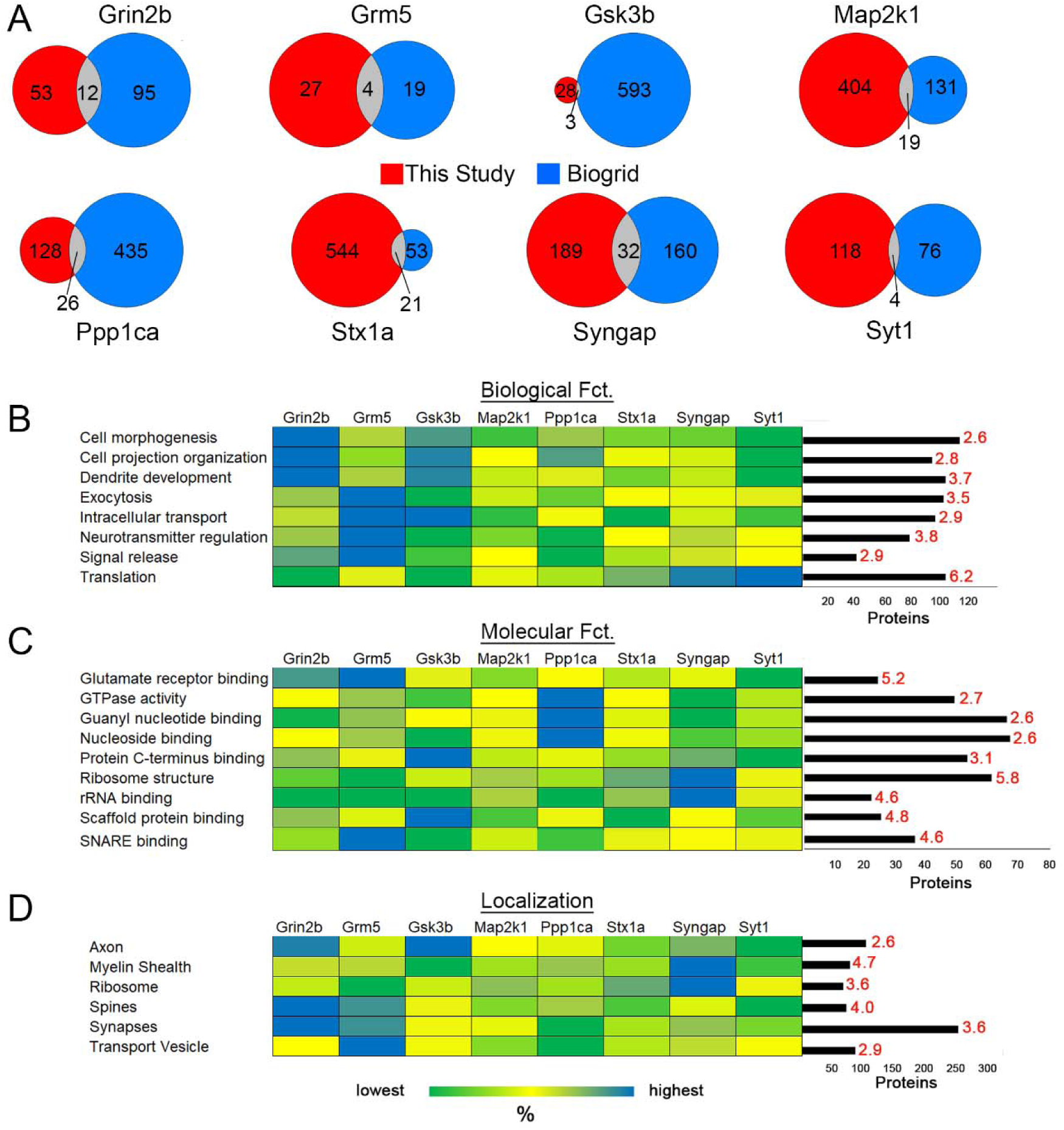
**A**, Comparison of the interactors of each bait in this study and BioGrid. Identification of significant GO enrichment terms of the SCZ disease network for three analyses: **B**, biological function (p < 2.22e^-16^), C, molecular function(p < 1.28E^-10^), and D, cellular component or localization (p < 2.2e^-16^). The bar graph depicts the number of interactors assigned to each enriched GO term. The red number above each bar represents the enrichment value over the background whole brain proteome. The heatmap depicts the percentage of interactors from each bait annotated to the enrichment GO term.

### Biological Interpretation of SCZ PPI network

The significant enrichment of GO terms was determined for the entire SCZ PPI network (**Figure2B**). The enriched biological functions included dendrite development, neurotransmitter regulation, intracellular transport, and translation. At the molecular level, glutamate receptor binding, SNARE binding, scaffolding binding, and ribosome structure were significantly enriched. The frequency of these functionally annotated proteins varied among the different baits. The Grm5 network had the highest percentage of proteins annotated to intracellular transport. This corresponds to reports that describe Grm5 as an atypical GPCR with more intracellular localization than plasma membrane localization [63, 64]. Syngap1 had the highest percentage of proteins involved in translation and ribosome structure regulation, which relates to studies demonstrating that it can regulate protein synthesis [65, 66]. Many different cellular components were also significantly enriched. Transport vesicles and ribosomes were enriched, consistent with the enriched biological functions. The cellular compartment with the most annotated proteins was the synapse. Synaptic proteins were observed in all the interactomes, but Grin2b interactome had the highest percentage of synaptic proteins. These synaptic proteins were validated and further investigated with the SynGo synaptic database [56]. This database mapped 41% (418 protein) of the interactors to unique synaptic genes, which was calculated as a significant enrichment (p-value = 2.6 e^-175^) (Figure 3A and ***Supplementary Table3***). In addition, 35 sub-synaptic child terms were significantly enriched with a 1% FDR. These annotated synaptic proteins were evenly distributed between the two largest sub-synaptic compartments, the pre- and post-synapse (Figure 3B). The SynGo database assigns a post-synaptic localization to all the baits except for Syt1 which was not designated to either compartment. Grin2b, Gsk3b, Ppp1ca, and Stx1a were also annotated to the pre-synapse. The proteins annotated exclusively to the pre- or post-synapse were evenly distributed among the bait’s SAINT interactors except Grin2b, Grm5, and Gsk3b which had more interactors assigned to the post-synapse (Figure 3C). Thus, this data demonstrates that many of these novel PPI may occur at the synapse. It has been reported that SCZ susceptibility proteins interact with one another beyond the level expected by chance[17]. The baits have been reported as SCZ susceptibility genes by several published studies analyzing large SCZ cohorts employing a variety of genetic methods, including GWAS, de novo mutation analysis, and differential methylation analysis. Cross-referencing these published genetic studies with our PPI network, 68% (684) of our protein interactors have evidence of being a potential SCZ risk factor. Within each bait analysis, most of the interactors have been implicated in SCZ susceptibility in previous studies (Figure 3D). Thus, our SCZ network corroborates the theory that SCZ susceptibility proteins form a physical interaction network.

**Figure 3.**
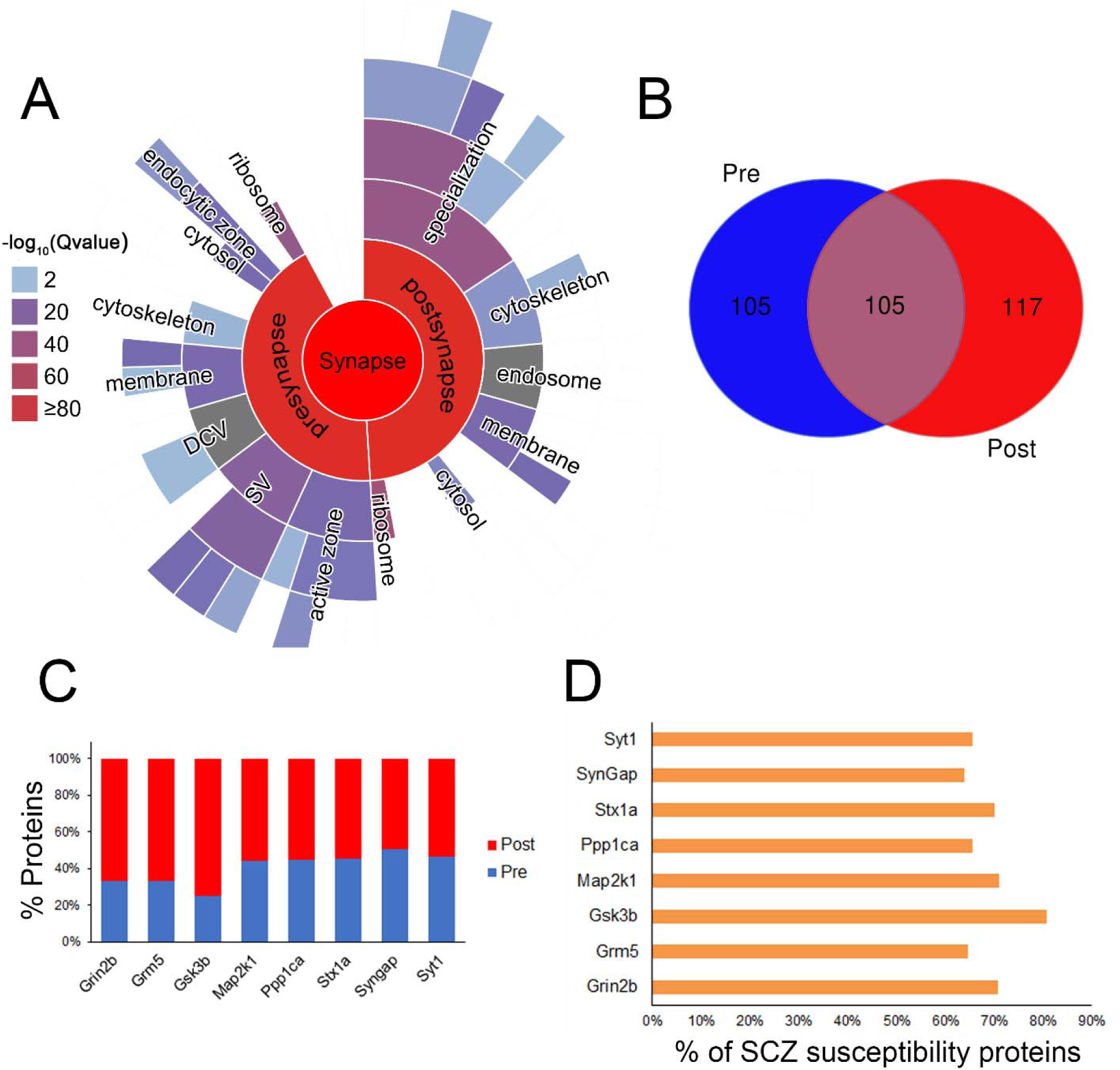
**A**, Significant enrichment of synaptic subcellular compartments. **B**, The overlap of pre- and post-synaptic proteins in A. **C**, The percentage of proteins exclusively pre- and post-synaptic proteins assigned to different baits. **D**, The percentage of SCZ susceptibility proteins assigned to each baits.

### APCP modulates the SCZ PPI network

We determined how PCP affected the identification of a highly confident PPI. First, the distribution of PPI between PCP and SAL networks were compared. Sixty-five percent of the PPI were only observed in SAL and only 12% were specific to PCP brains (Figure 4A). This suggests that PCP causes a widespread disruption of native PPI. However, this trend was not observed with all the baits (Figure 4C). The majority of PPI for Grin2b, Grm5, Gsk3b, and Syt1 were observed in PCP and not SAL, suggesting that PCP induces novel PPI for these proteins. Next, the SAINT datasets were re-examined with less stringent filtering (i.e., 20% FDR). On average, 36% of the high confidence PPI (i.e., 5% FDR) specific to one condition were observed in the other condition at lower confidence of 20% FDR (Figure 4B). This suggests that PCP may weaken or enhance some PPI rather than completely turning them “on” or “off”.

**Figure 4.**
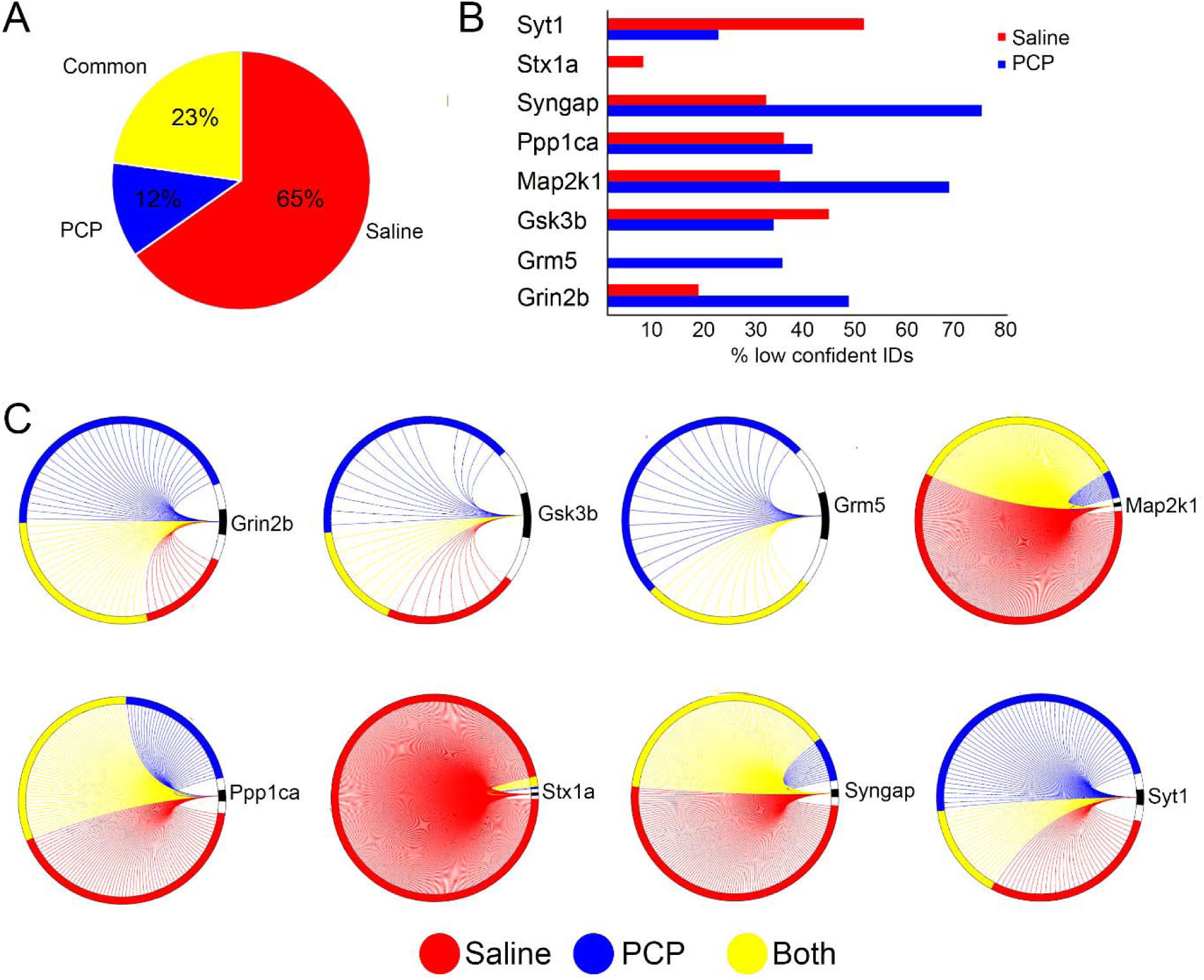
**A**, The pie graph represents the percentage of PPI identified by SAINT in PCP, saline, or both drug datasets for all the baits. **B**, The percentage of PPI with a 20% SAINT FDR in one drug condition and a 5% FDR in the other drug condition. **C**, Each circle represents one bait (black) and the interactors that were identified in PCP, Saline or both datasets by SAINT. Each line represents a unique interactor for that bait.

To further investigate the effect of PCP, the direct differences between SAL and PCP datasets for each bait were quantitated by employing the ^15^N internal standard[46]. Significant changes induced by PCP were observed for all baits except Syt1 (Figure 5A **and *Supplementary Table4***). The entire network revealed a 60% decrease of PPI upon PCP treatment. Grin2b, Gsk3b, and Map2k1 did not follow this trend, instead demonstrating most PPI increased with PCP. The enriched GO terms were like those seen in the SAINT analysis but included GO terms involved in synaptic biology (Figure 5B). Twenty-seven percent of the proteins significantly altered by PCP were also deemed a PPI by SAINT (FDR < 0.05) (Figure 5C). Interestingly, six significantly altered proteins are interactors with Gsk3b by the BioGrid database but not identified as a PPI with our SAINT analysis (Figure 5D). This suggests that adding a quantitative dimension to the PPI mapping aids in identifying interactors.

**Figure 5.**
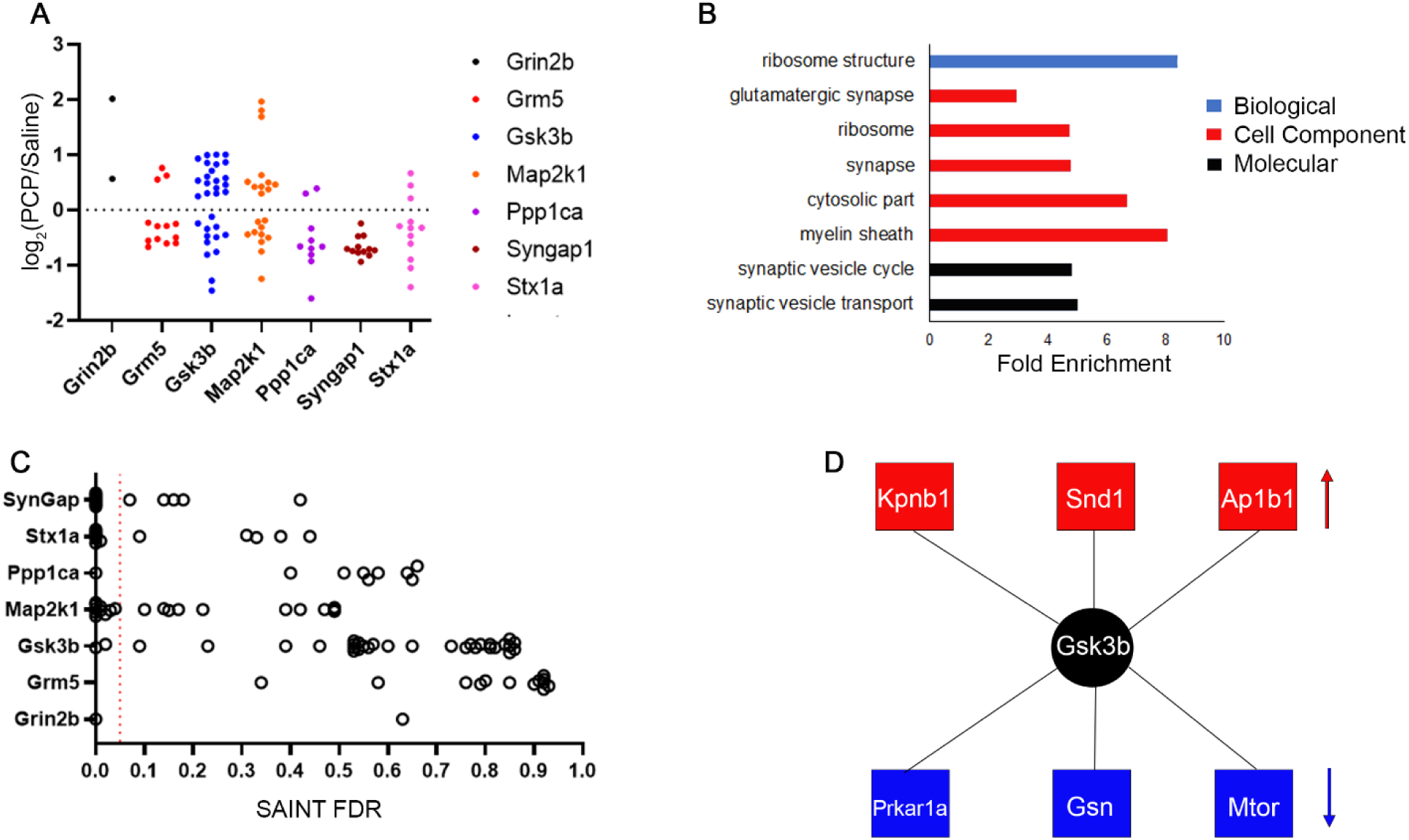
**A**, Altered proteins (p < 0.05) between PCP and SAL MS-IP datasets. **B**, Significantly enriched (p < 0.0002) GO terms. **C**, Altered proteins correlated with their SAINT FDR. Red dotted line indicates p-value 0.05. D, Gsk3b interactors the Biogrid database and significantly altered in the Gsk3b MS-IP experiments, but not assigned as a Gsk3b interactor b the SAINT analysis. Red indicates up-regulated and blue is down-regulated with PCP.

## Discussion

We performed *in vivo* PPI network mapping of eight genes that have been implicated as SCZ risk factors. Many previously reported PPI were identified, including ones needed for their canonical function. For example, Grin2a and Grin1 [67] were identified as Grin2b interactors because they form a calcium channel (i.e. NMDAR) with Grin2b.

Similarly, Ppp1ca is a catalytic subunit of the phosphatase PP1, which is required to associate with regulatory subunits to create holoenzymes [68]. Eleven different PP1 subunits were identified in the Ppp1ca interactome. Nevertheless, most PPI identified here are components of transient signaling complexes or have unknown functions. For example, our SCZ network validated the interaction between Grm5 and Homer, which directly binds the C-terminal tail of Grm5 [69]. Syngap1’s interaction with Fmr1 was validated in the study [30], but the function is not known. It has been reported that the signaling pathways of these proteins are functionally linked in the brain [65], providing evidence of a function interaction. Synaptopodin was validated as a Grm5 PPI and has been reported to regulate Grm5 membrane localization but is probably not a direct interaction [70]. Thus, our SCZ PPI network consists of two types of PPI: direct physical interactions and “co-complex” interactions. Our data cannot distinguish between these two interaction types unless previously reported or until further experiments are performed. Overall, this study validated 124 previously reported PPI, which is less than 10% of our PPI network. The large number of novel PPI most likely stems from the scarcity of MS-IP experiments previously performed in brain tissue. There were 688 publications in BioGrid where a PPI was reported with at least one of our baits and 22% (148) were high-throughput unbiased experiments. Only 9% (13) of these high-throughput experiments were performed in brain tissue and only one of these experiments shared a bait (i.e., Syngap1) with our study[71]. Correspondingly, our Syngap1 dataset did have the largest overlap with BioGrid. Overall, our data indicates that IP-MS studies in brain tissue have a huge potential to uncover novel PPI for a greater molecular understanding of neurobiology and to improve *in silico* network analyses for neurological diseases.

There were similarities between our *in vivo* SCZ PPI network and *in silico* SCZ genetic networks. First, *in silico* studies have demonstrated that SCZ risk factors are highly interconnected compared with proteins not implicated in SCZ. Our entire SCZ PPI network was interconnected with 42% of the 1007 interactors annotated as a highly confident interactor with one or more SCZ bait. There were many reciprocal PPI verified by two baits. Gsk3b and phosphatase PP1 have been reported to interact. Gsk3b was identified as a Ppp1ca interactor, and two PP1 subunits (Ppp1cc and Ppp1r2) were identified as interactors of Gsk3b. Similarly, members of the synaptotagmin protein family were identified as PPI to Stx1a and members of the syntaxin protein family were identified as PPI to Syt1 confirming their localization to a multi-protein complex essential for neurotransmission and vesicular fusion[72]. A Syngap1-Map2k1 interaction was originally reported with a MS-IP experiment with Syngap1 as the bait like this study but with a different Syngap1 antibody[71]. For the first time, this PPI was also identified with a Map2k1 antibody. The function of this reproducible PPI is unknown. This study also discovered a novel interaction between two baits: Grm5 and Stx1a. This PPI was identified in both the Grm5 and Stx1a datasets. Stx1a has been co-localized to Grin2B and Dlg4 at the hippocampus postsynaptic density, and it has been posited that Stx1a regulates the trafficking of Grin2b[73]. Grm5 is structurally linked to Grin2b via the scaffolding proteins Homer and Dlg4. Activation of Grm5 can potentiate NMDAR activity and NMDAR activation can desensitize Grm5[61, 74, 75]. This functional relationship has been proposed as therapeutic target for SCZ[76]. There were 11 interactors identified in both the Grm5 and Stx1a MS-IP datasets that are potentially involved in a Grm5-Stx1a complex. Interestingly, one is Homer1 and another is syntaxin family member Stx13, which has been reported as an interactor of Homer1[77]. This suggests that Stx1a-Grm5 may be involved in the functional relationship between NMDAR and Grm5.

Our PPI network was enriched (68%) in proteins previously associated with SCZ. For example, the neurexin (Nrxn) family are presynaptic proteins that regulate synaptogenesis and plasticity by binding postsynaptic cell adhesion proteins called neuroligins (Nlgn)[78]. Mutations in both neurexins and neuroligins are associated with SCZ[79–81]. Nrxn1 and Nrxn3 were identified as a PPI for Syt1 and Nlgn3 was identified as a PPI for Grin2b, Map2k1, and Stx1a. Nlgn1 and NMDAR are colocalized at postsynaptic density and modulation of Nlgn can regulate NMDAR function[82, 83].

Voltage-gated calcium channels (VGCCs) are enriched in SCZ risk factors [18, 84, 85], and our SCZ network included seven VGCCs. Similarly, six different subunits of voltage-gated potassium channels were observed in our network. Three (Kcna4, Kcna6, and Kcnab2) were Grin2b interactors and both Kcna4 and Kcnab2 are associated with SCZ. These channels can be homo- or heterotetrameric, so it is possible these three subunits form one functional potassium associated with NMDAR[86]. Kcna4 has been demonstrated to bind to the scaffolding proteins Dlg1 and Dlg4[87]. These scaffolds also bind Grin2b and were identified as Grin2b interactors in this study. This raises the possibility that these proteins are in a complex linked by Dlg4. Further research is needed to determine if a protein complex of NMDAR and a potassium channel modulate each other or if disruption of this complex may contribute to SCZ pathogenesis.

Our SCZ network was quantitated in response to PCP, which has been used to mimic a schizophrenic state in laboratory animals. Since the PCP treatment is less than 30 minutes, any alteration in transcription or translation can be ruled out, as that typically takes an hour to be triggered or detected[88, 89]. This suggests that the behavioral changes induced by PCP are mediated by alterations in post-translational modifications, protein trafficking, or PPIs. Indeed, using the protocol employed here, PCP has been reported to alter the phosphoproteome [48]. Phosphoproteins altered by PCP were enriched in similar pathways and functional groups observed in our SCZ PPI network including synaptic transmission and the cell projection organization. In this study, the effect of PCP was analyzed in two ways. First, the identification of PPI for each bait was analyzed with or without PCP using the SAINT algorithm. This analysis suggests that PCP weakened many PPI. For example, Grin2b binds Grin2a to form the well-studied functional NMDAR. Grin2a was observed as a PPI of Grin2b with PCP but not with SAL using a 5% FDR. It was identified as a Grin2b interactor with SAL at 20% FDR. Quantitation using ^15^N labeled brains (i.e., SILAM, Stable Isotope Labeling in Mammals) was also used to study the effects of PCP [90]. In the SAINT analysis, the PPI probabilities were compared between PCP and SAL datasets. In contrast, SILAM was used to directly quantitate proteins in the IPs without determining if a PPI exists, but the data still validated the SAINT results. For example, Dlg1 was observed as a Grin2b PPI with PCP, but not SAL. In the SILAM analysis, there was a significant increase of Dlg1 in the Grin2b IP upon treatment with PCP. Dlg1 binds directly to Grin2a and regulates the trafficking of Grin2A to the synapse[91]. NMDAR agonists increase the phosphorylation of Dlg1 which disrupts this interaction, thus reducing Grin2A insertion in the synaptic membrane. Consistent with this report, PCP has been demonstrated to reduce the phosphorylation of Dlg1[48]. Our data suggests that PCP decreases the Dlg1 phosphorylation and drives more Grin2A-Dlg1 to the post-synaptic membrane to form a complex with Grin2b. This hypothesis is also supported by our Grin2b network where Grin2a was identified as a highly confident interactor (5% FDR) with PCP whereas with SAL it was identified as a PPI but with a less robust signal (20% FDR).

The improper trafficking of NMDAR has been implicated in cognitive disorders[92]. The effects of PCP on our SCZ network suggests that disruption of NMDAR trafficking may also contribute to SCZ pathogenesis. Supporting this hypothesis, a recent study demonstrated altered NMDAR trafficking in three different models of SCZ[93].

## Conclusions

Although using MS for identification of PPI has been reported for decades, there still is scarce MS-IP studies in mammalian tissues. This data is essential to generate robust tissue specific *in silico* networks, which will allow the identification of disease “hubs” that could be potential drug targets. Currently, almost 40% of approved drugs target individual receptors, which include GPCR and ligand gated ion channels. Many receptor complexes are tissue or cell specific compared to the individual receptor expression, which allows differential receptor signaling in specific cell types. Thus, targeting complexes could lead to more selective drugs with fewer unwanted side effects[94].

Although this approach does have its own challenges, numerous PPI drugs have been tested in clinical trials, including drugs targeting the NMDAR-Dlg4 interaction[95]. The development of drugs targeting PPI is in its infancy, but it holds great promise for neurological disorders because brain regions possess discrete function and proteomes. The first step to this exciting drug pipeline is mapping PPI by proteomic studies as described here.

## Supporting information

SuppTable1

SuppTable2

SuppTable3

SuppTable4

SuppFig1

## Acknowledgements

This work has been supported by NIH grants 1R01 AG077046-01, R01 MH067880 (to J.R.Y.) and Veterans Affairs VISN 22 Mental Illness Research, Education and Clinical Center (MIRECC) (to S.B.P.) We thank Dr. Claire Delahunty(TSRI) for critical review of the manuscript.

## Author Contributions

Conceptualization, D.B.M and J.R.Y.; Investigation, D.B.M, S.B.P.; Formal Analysis and Visualization, D.B.M; Writing – Original Draft, D.B.M.; Writing – Review & Editing, D.B.M., S.B.P., and J.R.Y.; Funding Acquisition and Project Administration, J.R.Y. and S.B.P.; Supervision, J.R.Y

