## Supplementary material for "In vivo mapping of protein-protein interactions of schizophrenia risk factors generates an interconnected disease network": SuppFig1

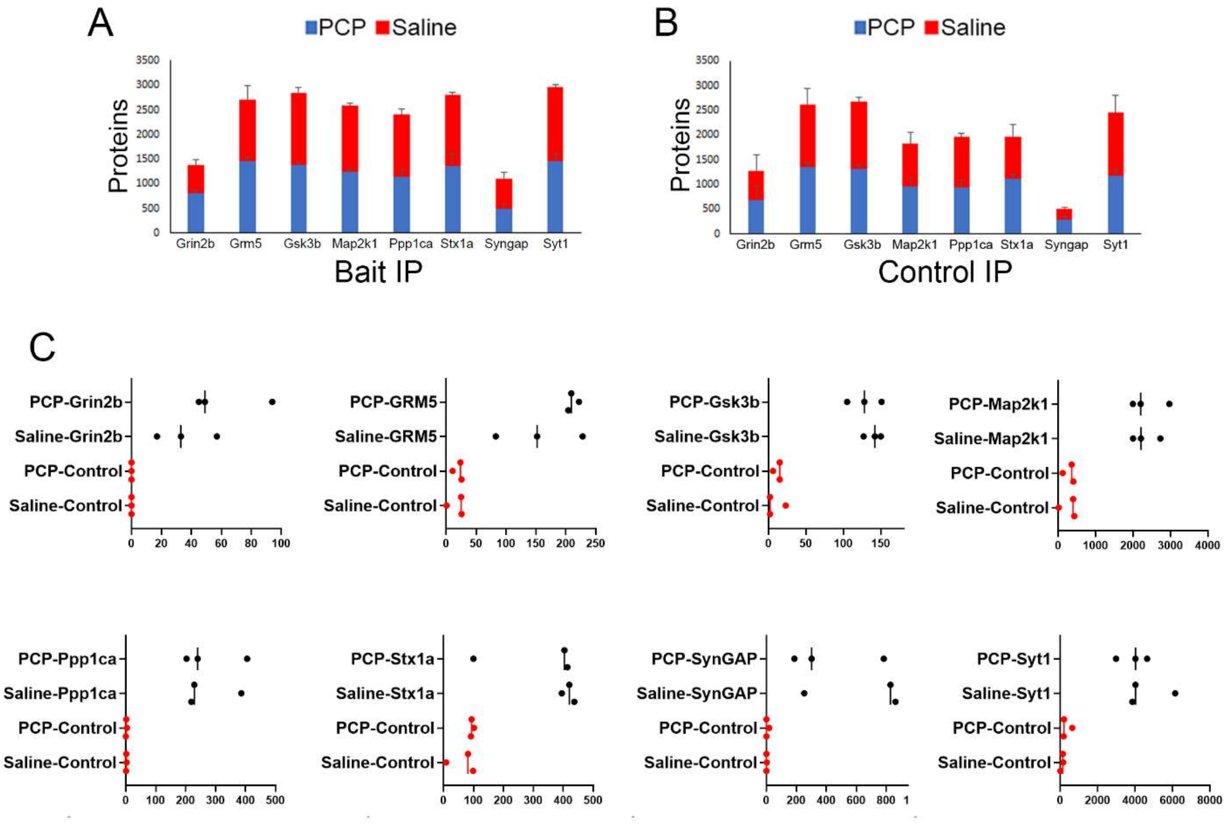


Supp. Figure 1

**Supplementary Figure 1.** The number of protein identified in the bait(**A**) and control(**B**) IPs. **C**, The abundance of the baits identified in the bait and control IPs. X-axis is spectral count.
